# Tubulin engineering by semisynthesis reveals that polyglutamylation directs detyrosination

**DOI:** 10.1101/2022.09.20.508649

**Authors:** Eduard Ebberink, Simon Fernandes, Georgios Hatzopoulos, Ninad Agashe, Nora Guidotti, Timothy M. Reichart, Luc Reymond, Marie-Claire Velluz, Fabian Schneider, Cédric Pourroy, Carsten Janke, Pierre Gönczy, Beat Fierz, Charlotte Aumeier

## Abstract

Microtubules, a critical component of the cytoskeleton, carry combinations of post-translational modifications (PTMs), which are critical for the regulation of key cellular processes. Long-lived microtubules, in neurons particularly, exhibit both detyrosination of α-tubulin as well as polyglutamylation. Dysregulation of these PTMs results in disease, including developmental defects and neurodegeneration. Despite their importance, the mechanisms governing the emergence of such PTM patterns are not well understood, mostly because tools to dissect the function and regulation of tubulin PTMs have been lacking. Here, we report a chemical method to produce fully functional tubulin carrying precisely defined PTMs within its C-terminal tail. Using a sortase- and intein-mediated tandem transamidation strategy, we ligate synthetic α-tubulin tails, which are site-specifically glutamylated to specific extents, to recombinant human tubulin heterodimers. Using microtubules reconstituted with such designer tubulins, we show that polyglutamylation of α-tubulin promotes its detyrosination by enhancing the activity of the tubulin tyrosine carboxypeptidase vasohibin/SVBP in a manner dependent on the length of polyglutamyl chains. Moreover, modulating polyglutamylation levels in cells results in corresponding changes in detyrosination. Together, using synthetic chemistry to produce tubulins carrying defined PTMs, we can directly link the detyrosination cycle to polyglutamylation, connecting two key regulatory systems that control tubulin function.

## Main

Microtubules are dynamic polymers composed of α/β tubulin heterodimers that form a dense network throughout the cell, which is critical for intracellular transport, cell division, motility and polarization^1^. Within this network, post-translational modifications (PTMs)of the tubulin heterodimers enable microtubules to adapt to specific cellular functionsJanke, 2020 #5255}. PTMs are particularly enriched on the unstructured tubulin C-terminal tails (CTTs). Here, PTMs such as polyglutamylation, i.e. the attachment of polyglutamate chains to specific glutamate residues within α- and β-tubulin, or detyrosination, i.e. the proteolytic removal of the C-terminal tyrosine residue of α-tubulin, coexist to form a combinatorial ‘tubulin code’, associated with distinct microtubule function^2,3^.

Specific ‘writer’ and ‘eraser’ enzyme classes exist for each PTM. The CTT of α- and β-tubulin CTTs is polyglutamylated by a family of Tubulin Tyrosine Ligase Like (TTLL) enzymes^4^, whereas glutamates are removed by the M14 carboxypeptidases family (CCPs) ^5^. Conversely, detyrosination is catalyzed by the carboxypeptidases vasohibin 1 and 2 (VASH1, VASH2), which exist in a heterodimeric complex with the small vasohibin-binding protein (SVBP)^6–9^. The levels of tubulin detyrosination and tyrosination, which itself is catalyzed by Tubulin Tyrosine Ligase (TTL), are critically implicated in cell function and development^10^. Moreover, both polyglutamylation and detyrosination are a hallmark of long-lived microtubules^11–15^, directly regulate microtubule stability^16^ and play major roles in regulating cellular transport processes^17–19^. Together, these PTM systems are of key importance for a plethora of cell functions, and their dysregulation is implicated in wide ranging disorders, most notably in developmental defects and neurodegeneration^20^. Thus, a detailed understanding of the molecular function of tubulin PTMs is of key importance. However, due to the complexity of the modifications, as well as the absence of methods to prepare tubulins containing defined PTMs, a systematic exploration of PTM functions and their inter-dependence is still lacking.

Direct isolation or purification of tubulin from natural sources (i.e. bovine brain) provides a complex mixture of differently modified tubulin heterodimers, which renders an investigation of individual PTM functions impossible. To directly dissect fundamental regulatory mechanisms governing the establishment of the tubulin code, a method to produce homogeneous microtubules with precisely installed and chemically defined PTMs is critically required. Earlier methods of controlling the modification state of tubulin, including the state of polyglutamylation, used either enzymatic modification, which however resulted in broad length distributions and unknown attachment points^16^. Alternatively, polyglutamate branches had been introduced via chemical modification of cysteine residues, albeit at low reaction yield and within the context of hybrid yeast/human tubulins^21^. Moreover, these methods were only applicable to a single type of PTM and were thus constrained in their general applications.

Here, we report the development of a synthetic approach that allows us to introduce any combination of PTMs at defined positions into tubulin CTTs with full chemical control. These semisynthetic designer tubulins represent novel tools which enable a molecular dissection of the tubulin code with high specificity. In the following, we then used these new reagents to demonstrate a novel crosstalk mechanism between two key tubulin PTM classes: We show that polyglutamylation of microtubules is required for the detyrosination activity of vasohibin/SVBP. In agreement, we find that modulating polyglutamylation levels in cells directly controls detyrosination. Taken together, our novel designer tubulins thus not only establish an intimate link between these two regulatory PTM systems, but also set the stage for a molecular dissection of the tubulin code.

### A semisynthetic strategy to produce modified tubulin

To introduce any combination of PTMs into tubulin with perfect regioselectivity, we decided to develop a protein semi-synthesis approach that relies on the ligation of a chemically synthesized CTT peptide, harboring PTMs, to recombinantly produced tubulin dimers.

Tubulin dimers are large (100 kDa) and sensitive enzyme complexes, which are difficult to express, to purify^22^ and quickly lose activity when exposed to non-native solvent conditions. To preserve the full functionality of tubulin, we used split-intein-based protein trans-splicing^23^, which is compatible with native solvent conditions and rapidly links synthetic and recombinant protein parts. Split-inteins consist of two fragments (Int^N^ and Int^C^) that fold upon contact and catalyze the connection of flanking protein sequences via a native peptide bond, splicing themselves out of the sequence in this process. For the implementation of this approach, we decided to split the tubulin sequence into two parts (**Figure 1a**): The N-terminal globular part of α-tubulin (amino acids 1-439), which is recombinantly expressed, and the tubulin CTT (amino acids 440-451), carrying the desired PTMs, which is chemically synthesized.

**Figure 1.**
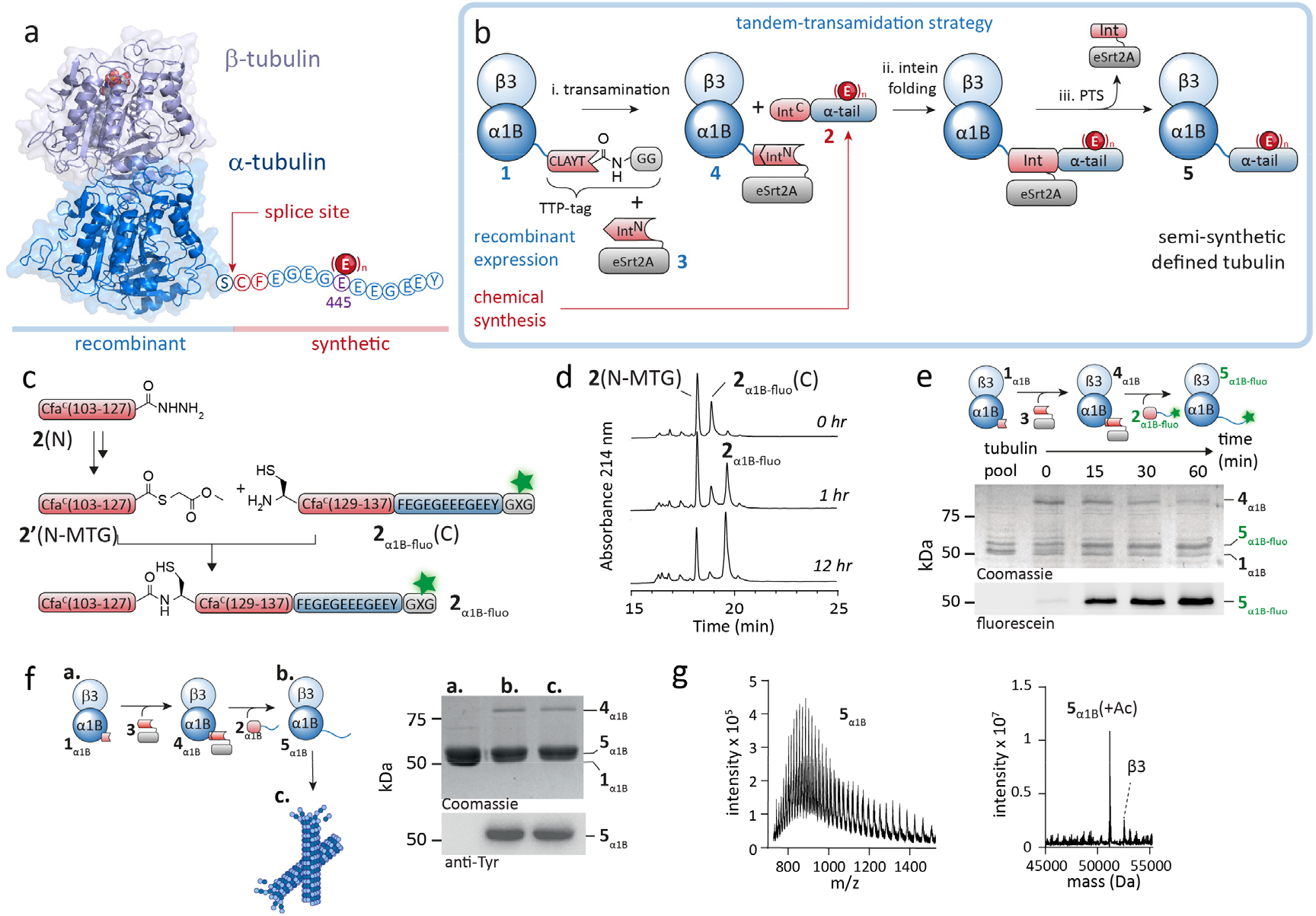
Semisynthetic tubulins to probe the tubulin code. **a**) Structure of tubulin heterodimer (PDB code: 1TUB) showing attachment site of the α-tubulin C-terminal tail (CTT). Indicated is the site of polyglutamylation E445, (E)_n_, as well as the splice site, with indicated sequence changes. **b**) Tandem-transamidation strategy to produce semisynthetic tubulin. Constructs with blue numbers were recombinantly expressed, construct with red number was chemically synthesized, see text for details. **c**) Synthesis of peptide **2**_α1B-fluo_ from two synthetic segments via NCL. X denotes homopropargylglycine, used for fluorophore attachment. For peptide synthesis and analytics, see **Supplementary Fig. 3**. **d**) RP-HPLC analysis of ligation reaction between **2**(N) and **2**_α1B-fluo_(C) to produce **2**_α1B-fluo_. **e**) SDS-PAGE analysis, visualized by Coomassie blue (top) and fluorescein fluorescence emission (bottom) of the tandem-transamidation reaction between **1**_α1B_ and **2**_α1B-fluo_ yielding the fluorescent α-tubulin **5**_α1B-fluo_. **f**) Synthesis reaction of **5**_α1B_, followed by polymerization into microtubules. Microtubule (pellet) and tubulin (supernatant) were separated by ultracentrifugation. Polymerization was monitored by SDS-PAGE and visualized by Coomassie blue (top) and a tyrosine specific antibody (bottom). Lanes: a, **1**_α1B_; b, reaction product **5**_α1B-_dimer supernatant; c, **5**_α1B_-microtubule pellet. See also **Supplementary Fig. 4c-e**. **g**) Liquid-chromatography mass spectrometry (LC-MS) analysis of ligated α-tubulin **5**_α1B_. Left panel: m/z spectrum, right panel: Deconvolved spectrum, showing the expected mass of **5**_α1B_ + 1Ac (calc: 51169 Da, obs.: 51168 Da).

In initial experiments we realized that attaching the large Int^N^ fragment to the α-tubulin C-terminus resulted in tubulin misfolding and inactivation (**Supplementary Fig. 1a**). We therefore used a tandem-transamidation reaction cascade, based on transpeptidase-assisted intein ligation (TAIL)^24^, to connect the synthetic tail peptide to the recombinant α-tubulin (**Fig. 1b**). Here, the C-terminal tail of α-tubulin is replaced by a short peptide sequence, the tandem-transamidation peptide tag (TTP-tag), resulting in tubulin construct **1**. In addition, a modified CTT is synthesized (peptide **2**), carrying the short C-terminal half of the engineered consensus fast (Cfa) DnaE split-intein^25,26^ (IntC) (**Fig. 1b** and below). In a first step, the N-terminal half of Cfa (Int^N^) is then connected to α-tubulin via a transamidase, the engineered bacterial sortase eSrt2A^27,28^ (**Fig. 1b**, step i). Fusing Int^N^ to sortase (Int^N^-eSrt2A, **3**) improves both yield and suppresses side-reactions in this step^24^. After the attachment of Int^N^ to tubulin, the Int^C^-CTT peptide **2** is added, resulting in intein reconstitution (**Fig. 1b**, step ii) followed by protein trans-splicing and the production of semisynthetic tubulin (**Fig. 1b**, step iii). The obtained tubulin sequence is almost native except for the insertion of a cysteine at the splice-site that is required for intein activity and an amino acid change (Val441 to Phe) for improved splicing efficiency (**Fig. 1a**).

### Semisynthesis of functional tubulin heterodimers

We first validated that the TTP-tag did not disturb α1B tubulin function by overexpressing the construct in Hela cells (**Supplementary Fig. 1a**). We then recombinantly expressed the tubulin heterodimer α1B/β3 (denoted as **1**_α1b_) in insect cells (**Supplementary Fig. 1b,c**) ^22,29,30^. For purification, we added a C-terminal FLAG-tag to β3, cleavable by TEV protease for optional removal^29,31^, and inserted an internal His6-tag into an internal loop of α1B^31,32^. This allowed us to obtain tubulin heterodimers **1**_α1B_ from insect cell culture in milligram amounts (2-3 mg per 600 mL insect cell culture) and high purity (**Supplementary Fig. S1d,e**).

For tubulin semisynthesis, we further required peptide **2**, corresponding to an appropriately modified tubulin CTT fused to the Cfa IntC sequence (Cfa^C^) (**Fig. 1a,b**). In an initial step to optimize the splicing reaction conditions, we designed a peptide corresponding to IntC followed by a α1B CTT without any PTMs, but carrying a fluorescent dye for facilitated detection of the reaction product. Moreover, to increase modularity and improve peptide synthesis yield, we divided peptide **2** into two parts that were separately synthesized (**2**(N) and **2**_α1B-fluo_(C), **Fig. 1c** and **Supplementary Fig. 2**). The two peptides were subsequently ligated via native chemical ligation (NCL)^33^ in the presence of methyl thiolglycolate (MTG), to form peptide **2**_α1B-fluo_ within 12 hours (**Fig. 1d**, **Supplementary Fig. 2**). We then proceeded to connect the CTT peptide **2**_α1B-fluo_ with recombinant tubulin, following the general tandem-transamidation strategy outlined in **Fig. 1f** and **Supplementary Fig. 3a**. Upon addition of Cfa^N^-eSrt2A (**3**) (**Supplementary Fig. 1f**) to tubulin heterodimers **1**_α1B_, the intermediate **4**_α1B_ was formed within seconds (**Fig. 1e**). Following systematic optimization of the conditions, further reaction with CTT peptide **2**_α1B-fluo_ resulted in the formation of fluorescent tubulin heterodimers **5**_α1B-fluo_ over the course of 60 min (**Fig. 1e** and **Supplementary Fig. 3b**), with final yield of ~ 70%. The thus generated semisynthetic tubulin heterodimers **5**_α1B-fluo_ could polymerize into microtubules, validating that they are functional (**Supplementary Fig. 3c-g**).

To provide control conditions for tubulin code exploration, both semisynthetic unmodified and PTM-containing α1B/β3 tubulin heterodimers are necessary. We thus repeated the synthesis as described before using a CTT peptide containing solely the native α-tubulin CTT without a fluorescent label (**2**_α1B_) and thereby produced unmodified tubulin heterodimers **5**_α1B_ (**Supplementary Fig. 4-6**). We monitored both synthesis and subsequent microtubule polymerization via immunoblotting using an antibody specific for the terminal tyrosine in α1B (anti-Tyr) (**Fig. 1f** and **Supplementary Fig. 5c**), as well as mass spectrometry, showing conversion to **5**_α1B_ (**Fig. 2g** and **Supplementary Fig. 6**).

**Fig. 2.**
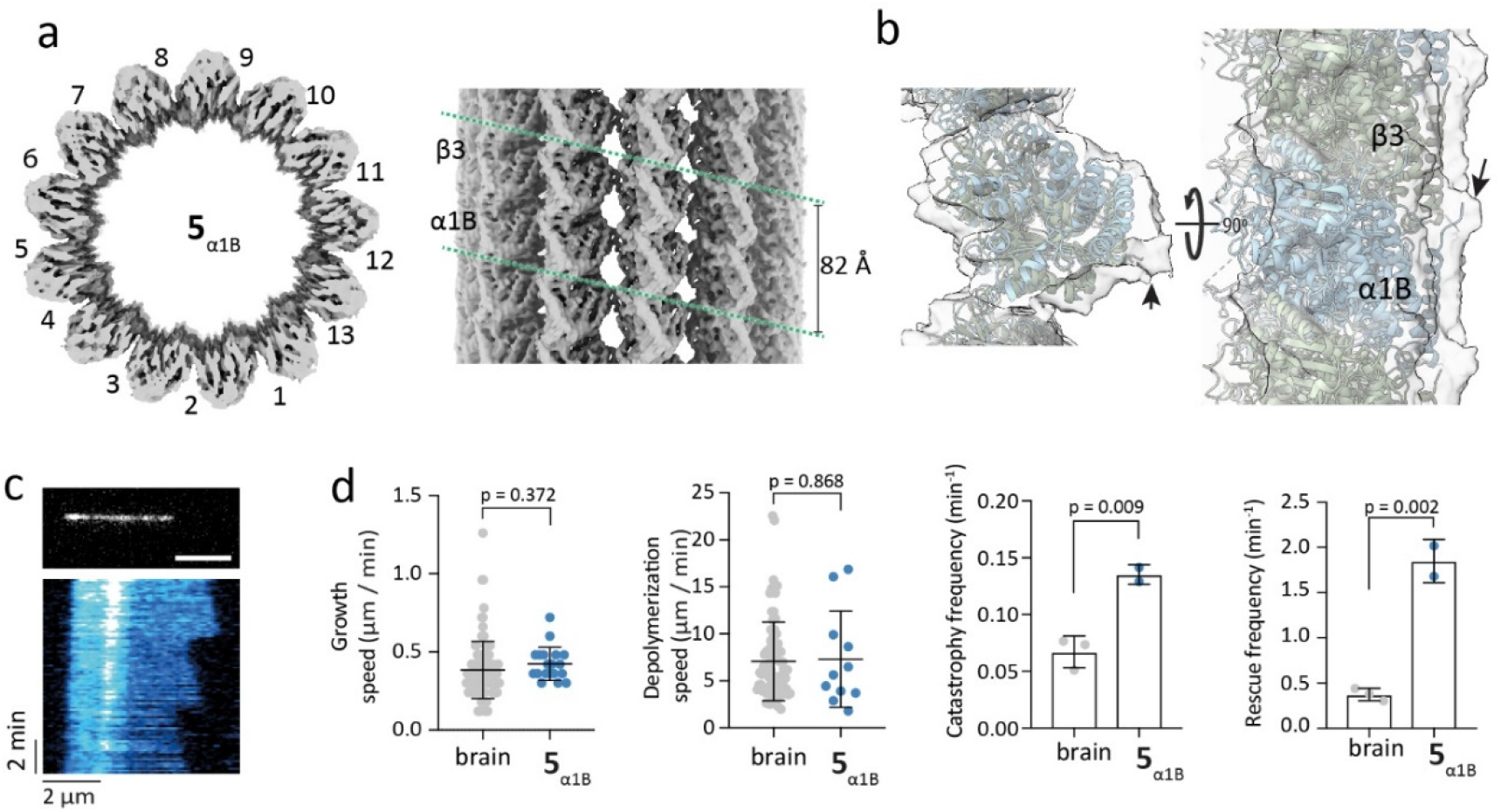
Structure and dynamics of semisynthetic microtubules. **a**) Top view (left) and side view (right) of cryoEM structure of taxol-stabilized GTP-microtubules polymerized from **5**_α1B_. **5**_α1B_ polymerized into a 13-protofilaments 3-start lattice. For the structure of **1**_α1B_ and **5**_α1B-5E_ microtubules see **Fig. S9**. **b**) A Cryo-EM map at lower threshold levels of for **5**_α1B_ shows additional density at the region of the CTT (arrow). **c**) Top: **5**_α1B_ microtubule visualized using 1 nM SiR-tubulin^35^. Bottom: Kymograph of **5**_α1B_ microtubule dynamics. **d**) Growth and shrinking dynamics, catastrophe and rescue frequency of **5**_α1B_ microtubules compared to bovine brain microtubules. Error bars: SD, Independent replicates n = 2 (**5**_α1b_), n = 3 (brain). Statistics: Student’s t-test.

To interrogate the structural integrity of semisynthetic microtubules, we determined their structure with cryo-EM, solving their structure to ~3.8 Å resolution (**Fig. 2a,b** and **Supplementary Figure 7**). This analysis revealed an organization into a 13 protofilaments 3-start lattice with an inter-dimer spacing of 8.2 nm, in line with previous reports from microtubules polymerized from bovine brain tubulin under similar conditions^34^. In addition, at lower threshold the EM map revealed further densities in α-tubulin tail regions, indicating the ligated tail domains; however, these were not well resolved, as anticipated from the likely flexible nature of the tails (**Fig. 2b**).

We further probed microtubule polymerization and depolymerization dynamics with TIRF-microscopy. Microtubules polymerized from **5**_α1B_ showed characteristic dynamic instability behavior, with successive growing and shrinking phases (**Fig. 2c**). The dynamic parameters, growth and shrinking velocities closely resembled the dynamic behavior of microtubules polymerized from bovine brain tubulin (**Fig. 2d**); however, unmodified semisynthetic tubulins showed a 2-fold increased catastrophe and 4-fold increased rescue frequencies (**Fig. 2d**). Our observation confirms previous reports that the β3-tubulin isotype has a destabilizing effect on microtubules, increasing the catastrophe frequency, while the impact on growth and shrinking velocities is minor^36–38^. In conclusion, tubulin heterodimers produced via our semisynthetic protocol are fully functional and retain all their dynamic and structural properties, thus enabling detailed investigations of the molecular functioning of the tubulin code.

### Detyrosination of tubulin is linked to its polyglutamylation

Microtubules often contain complex combinations of PTMs^5^. A possible mechanism underlying the establishment of defined PTM patterns is crosstalk between modifications^39^, where pre-existing PTMs influence writer and eraser enzyme activity, thereby facilitating or preventing the installation of additional PTMs. To investigate if the emergence of PTM patterns are dependent on PTM crosstalk, we analyzed the subcellular distribution of polyglutamylation and detyrosination in RPE-1 cells. Both modifications had a punctuated pattern along microtubules^40^ (**Fig. 3a** and **Supplementary Fig. 8c,d**). Polyglutamylation and detyrosination showed strong colocalization, beyond the probability of two individual signals to randomly overlap (**Fig. 3b,c** and **Supplementary Methods**). A majority of detyrosinated regions exhibited polyglutamylation (Mander’s overlap coefficient ΔY/pE, M = 0.63) and polyglutamylated regions were similarly associated with detyrosination (pE/ΔY, M = 0.37, Fig. **3c** and **Supplementary Fig. 8e**).

**Fig. 3.**
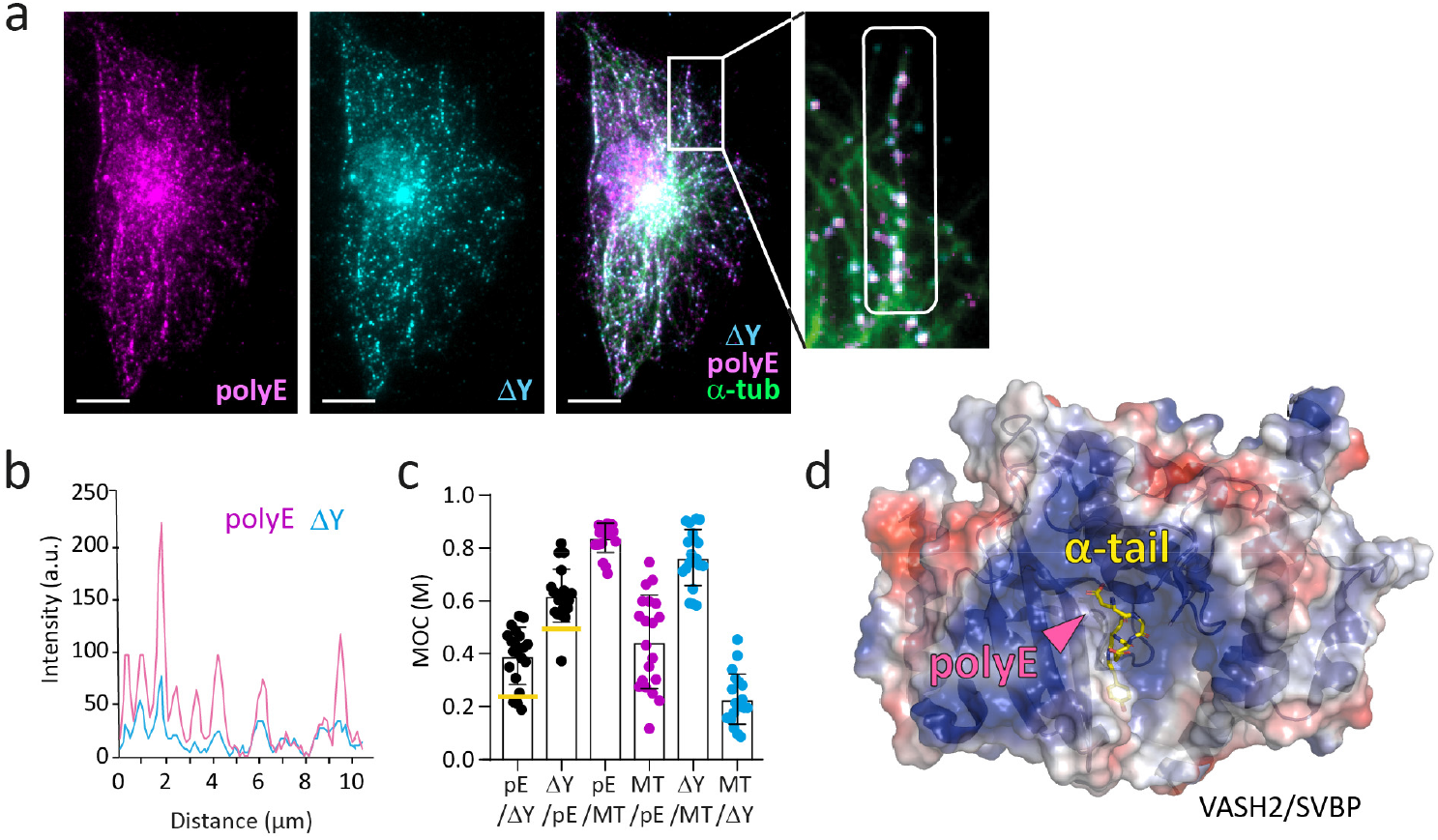
Polyglutamylation and detyrosination are colocalized in cells. **a**) Representative immunostaining of CRISPR/Cas9 knock-in GFP-Tubulin RPE-1 cells (green), antibody stained for polyglutamylation (polyE, magenta) and detyrosination (ΔY, cyan). The marked area in the inset shows a single microtubule. See also **Supplementary Fig. 8c**. Scalebar: 10 μm. **b**) Linescan along microtubule of the marked area in **a**). **c**) Colocalization of polyglutamylation (pE), detyrosination (ΔY) with microtubules (MT) analyzed via Mander’s overlap coefficient (MOC). The observed values for M(pE/ΔY) and M(ΔY/pE) indicate that the two modifications are not randomly distributed (yellow bar,Methods). **d**) Structure and surface electrostatics (red: negative, blue: positive) of vasohibin/SVBP (PDB code: 6J4V) bound to the α-tubulin tail (yellow). Magenta arrowhead indicates approximated position of polyglutamylation.

Based on these findings, as well as on the fact that polyglutamylation levels were higher than detyrosination in four different cell lines (**Supplementary Fig. 8a-b**), we hypothesized that the two different modifications are linked and polyglutamylation primes microtubules for detyrosination. We reasoned that the structures of vasohibin/SVBP, the enzyme responsible for tubulin detyrosination, provide a possible mechanism^7,8,41^. The vasohibin/SVBP heterodimer exhibits a positively charged surface surrounding its active site, where it binds the negatively charged CTT. Increased negative charge on the tubulin tails following polyglutamylation could thus modulate this interaction (**Fig. 3d**). We thus decided to test this hypothesis using semi-synthetic polyglutamylated microtubules.

### Assembling polyglutamylated tubulins

Polyglutamylation is a heterogeneous PTM with regards to polyglutamate branch length and attachment residues, which renders its functions difficult to investigate without chemical control. We thus set out to synthesize tubulin heterodimers carrying defined polyglutamate modifications within the α-tubulin CTT, allowing us to directly assess vasohibin/SVBP activity on differently polyglutamylated microtubules. For the branch point of glutamate attachment, we chose glutamate 445 (E445, **Fig. 1a**), as this site is most prominently polyglutamylated in neurons^5,42^. Following our modular strategy, we first synthesized a set of tubulin tail peptides, containing a branch of either five (E5) or ten (E10) glutamates. For the attachment of such polyglutamyl-branches, we replaced E445 by homopropargyl-glycine and used a Cu(I)-catalyzed azide-alkyne cycloaddition (CuAAC) coupling strategy^43^ resulting in a 1,2,3-triazole, a commonly used peptidomimetic linkage whose structural and electronic characteristics are similar to those of a isopeptide bond^44,45^ (**Fig. 4a** and **Supplementary Fig. 4b**).

**Fig. 4.**
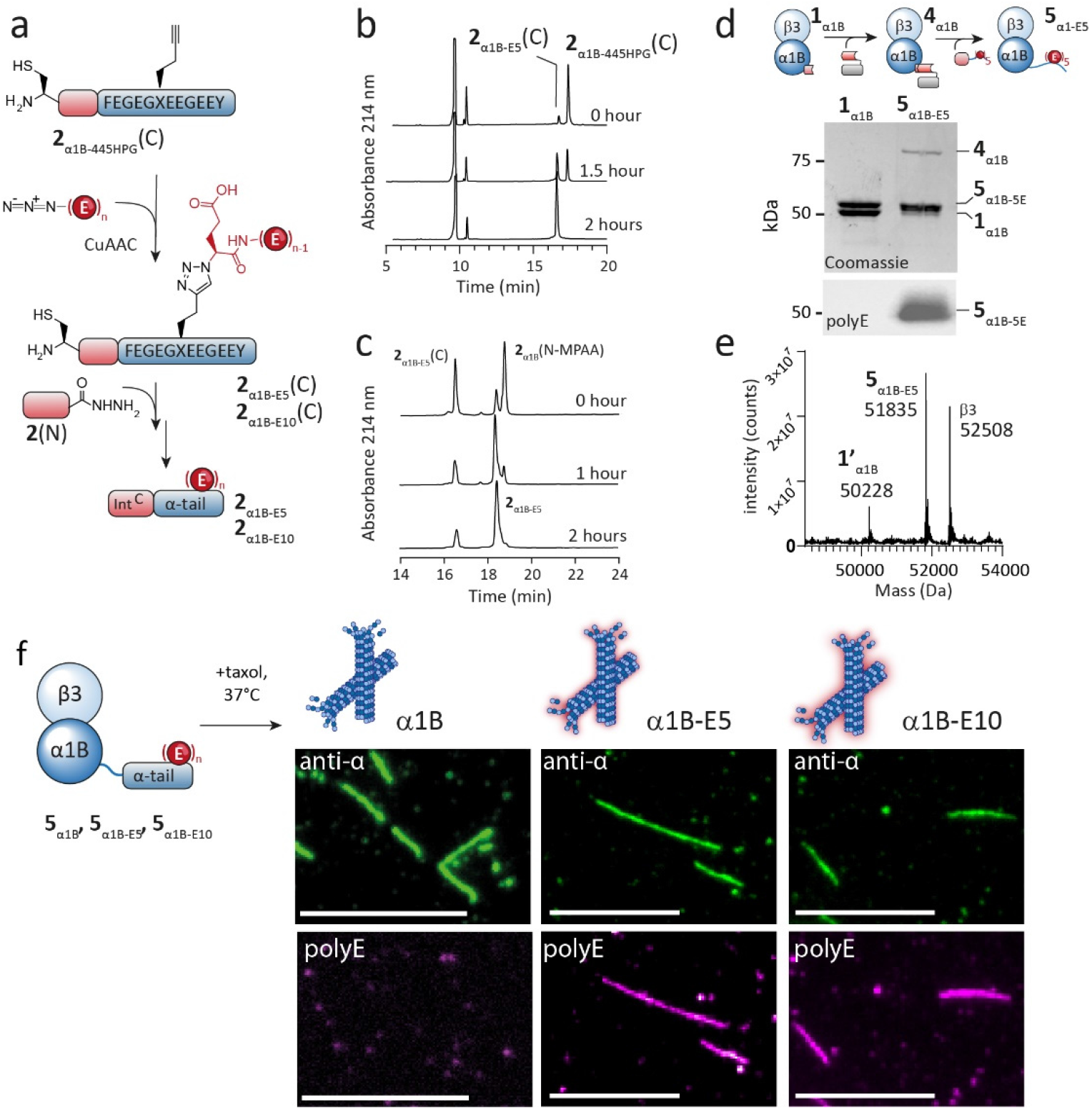
Synthetic polyglutamylated microtubules to probe crosstalk between polyglutamylation and detyrosinatione. **a**) Synthesis of polyglutamylated peptides **2**_α1B-E5_ and **2**_α1B-E10_ via CuAAC. **b**) RP-HPLC analysis of CuAAC reaction to synthesize **2**_α1B-E5_(C). **c**) RP-HPLC analysis of the ligation between **2**_α1B-E5_(C) and **2**(N) yielding **2**_α1B-E5_ within 2 hours. For analytical data, see **Supplementary Fig. 4**. **d**) SDS-PAGE and WB (anti-polyE) analysis of the semisynthesis of **5**_α1B-E5_. **e**) Deconvoluted mass spectrum of **5**_α1B-E5_/β3. **1’**_α1B_ corresponds to **1**_α1B_, lacking the final (Gly)2, and represents an unreactive side product. **f**) Generation of semisynthetic microtubules via polymerization of **5**α1B, **5**_α1B-E5_ and **5**_α1B-E10_. Microtubules are stained with indicated antibodies. Scalebar: 5 μm. For larger fields of views, see **Supplementary Fig. 10**.

We synthesized an azide-modified glutamate derivative (5-tert-butyl hydrogen 2-azidoglutarate, **Supplementary Methods**), which was included as the N-terminal amino acid in E5 and E10 chains (**Supplementary Figure 9a**). We prepared a tubulin CTT peptide containing homopropargyl-glycine at position 445 and coupled the polyglutamyl chains via CuAAC (**Fig. 4a,b**), followed by NCL to **2**(N) to generate peptides **2**_α1B-E5_ and **2**_α1B-E10_ (**Fig. 4c** and **Supplementary Fig. 9a**). With this set of modified peptides in hand, we proceeded to synthesize modified tubulin heterodimers **5**_α1B-E5_ and **5**_α1B-E10_ using the strategy delineated above (**Fig. 4d,e** and **Supplementary Fig. 4-6, 9**), allowing us to polymerize microtubules of defined modification states (semi-synthetic unmodified α1B as well as glutamylated α1B-E5 and α1B-E10).

We visualized the three different microtubules sedimented onto glass slides with TIRF microscopy, revealing their modified status with polyE-specific antibodies (**Fig. 4f, Supplementary Fig. 10**). Moreover, the cryo-EM structure of α1B-E10 microtubules was comparable to structural parameters of unmodified α1B and tail-less microtubules **1**_α1B_ (**Supplementary Fig. 7**). Interestingly, lower threshold EM-maps of α1B-E10 microtubules exhibited densities that could reflect the additional polyglutamylation (**Supplementary Fig. 7d-f**), albeit at low resolution. Finally, mono-glutamate branches could also be achieved directly via peptide synthesis, resulting in the native isopeptide bond (**Supplementary Figure 4c, 5c, 6b, 9c, 10b**), demonstrating that fully native PTMs can be prepared, if required.

Concluding, we developed a method to engineer functional, semisynthetic microtubules of desired type, composition and PTM position within the CTT. Such defined modified tubulin heterodimers now serve as novel tools to systematically dissect the effect of tubulin PTMs, their regulatory crosstalk and functional consequences.

### Revealing crosstalk between polyglutamylation and detyrosination

With specifically modified tubulins in hand, we then set out to determine if polyglutamylation alters the enzymatic efficiency of vasohibin/SVBP in removing the final tyrosine from the α-tubulin CTT. It has been shown that vasohibin/SVBP is not active on tubulin CTTs alone, but requires microtubules as a substrate^46^. We used α1B, α1B-E5 and α1B-E10 microtubules to determine vasohibin/SVBP-mediated detyrosination (**Fig. 5a**). We thus recombinantly expressed and purified vasohibin/SVBP (**Supplementary Fig. 11**), and incubated the complex with our three different semisynthetic microtubules carrying either an unmodified, 5E or 10E CTT. Co-pelleting assays did not reveal any significant differences of vasohibin/SVBP binding to α1B and α1B-E10 microtubules (**Fig. 5b** and **Supplementary Fig. 12**). In contrast, vasohibin/SVBP catalytic activity was highly sensitive to the polyglutamylation-state of the substrate microtubules: In the absence of polyglutamylation, vasohibin/SVBP showed very low activity, with barely any detectable detyrosination within 30 minutes (**Fig. 5c, d**).

**Fig. 5.**
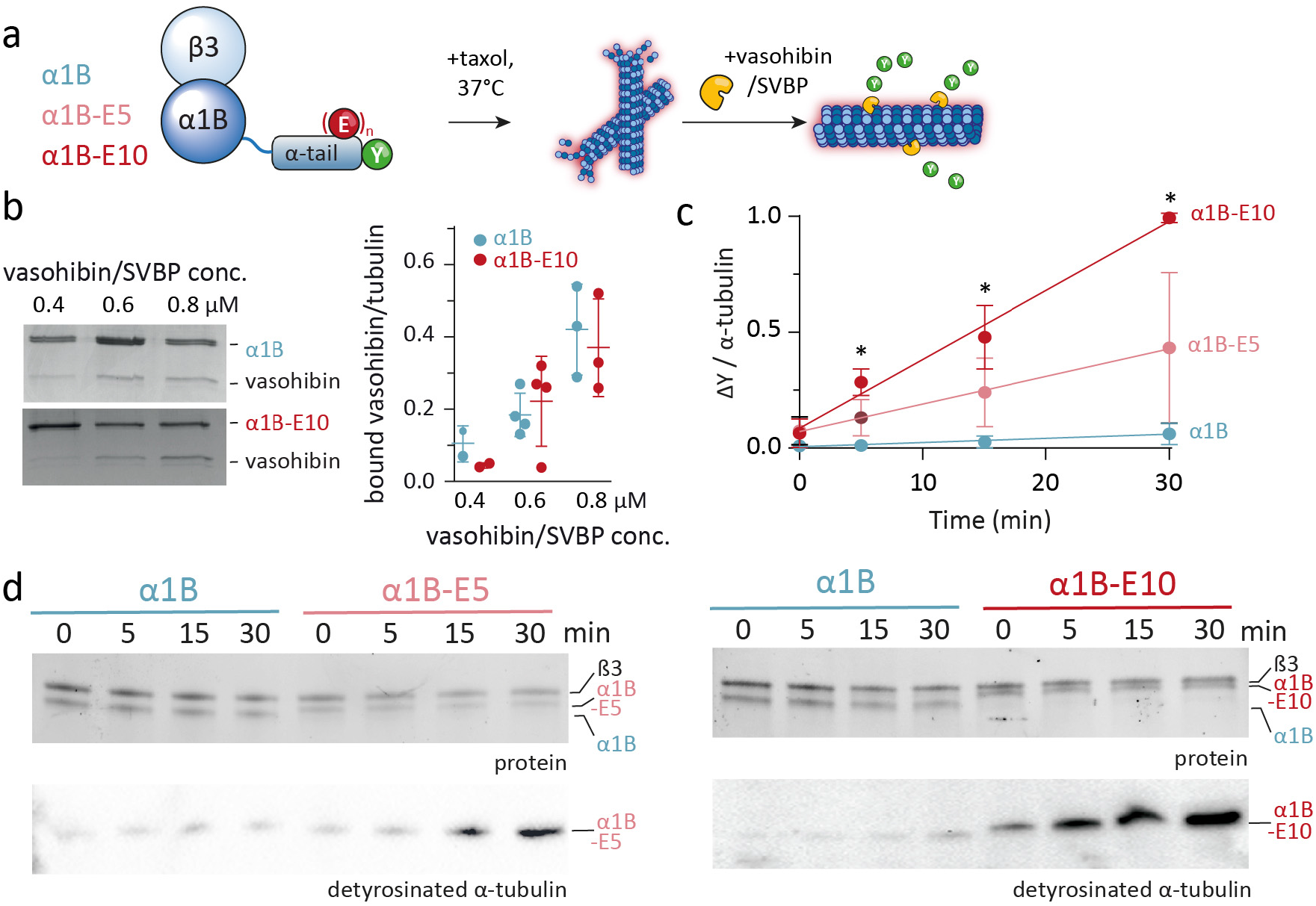
Tubulin polyglutamylation stimulates its own detyrosination by vasohibin. **a**) Schematic representation of the detyrosination assay. **b**) Percentage of vasohibin/SVBP at different concentrations bound to unmodified (blue) and polyglutamylated (red) microtubules (see **Supplementary Fig. 12**). Mean with SD, n = 2-4 independent experiments. Student’s t-test p > 0.05 for all conditions. **c**) Quantification of ΔY reactions. Detyrosination increases with the polyglutamylation level. Mean with SD, *: Student’t t-test, p<0.05; for all individual repeats and statistical analysis see **Supplementary 13d**. **d**) Western blot analysis of detyrosination by vasohibin/SVBP of α1B α1B-E5 and α1B-E10 using antibodies against ΔY α-tubulin.

Strikingly, the presence of polyglutamylation of the CTT stimulated vasohibin/SVBP activity. For α1B-E5 microtubules, detyrosination increased throughout the reaction time reaching up to 9-fold higher detyrosination levels compared to unmodified microtubules (**Fig. 5c,d** and **Supplementary Fig. S13a-d**). For α1B-E10, the effect was even more pronounced (**Fig. 5d**), resulting in a 16-fold increased vasohibin/SVBP detyrosination activity over unmodified microtubules (**Fig. 5c**). Quantifying the survival-fraction of tyrosinated α-tubulin revealed that after 30 min, α1B-E10 was substantially detyrosinated, in contrast to unmodified α1B (**Supplementary Fig. 13e**). In conclusion the level of polyglutamylation is not critical for vasohibin/SVBP recruitment, but increases its enzymatic efficiency.

### Crosstalk between polyglutamylation and detyrosination in cells

Does glutamylated tubulin act as a preferred substrate for detyrosination also in the complex cellular environment? To address this question, we overexpressed TTLL5 and TTLL6 enzymes adding mono- and polyglutamates to the CTT of tubulin^40^, in RPE-1 and U2OS cells. In RPE-1 cells, this resulted in an over 7-fold increase for both polyglutamylation and detyrosination (**Fig. 6a, Supplementary Figure 14a**), compared to the previously observed isolated PTM patches (**Fig. 3a**). Accordingly, a majority of polyglutamylated regions now showed detyrosination (pE/ΔY, M = 0.65, **Fig. 6b** and **Supplementary Fig. 14b**). This finding was confirmed using immunoblot analysis of cell lysates that showed a 13-fold increase in α-tubulin polyglutamylation levels, and a corresponding 7.1-fold increase of detyrosination (**Fig. 6c,d** and **Supplementary Fig. 14c**). Conversely, treating cells with a cocktail of siRNAs against five different TTLLs reduced the polyglutamylation level by 67 %, resulting in a concomitant reduction in detyrosination by 68 % (**Fig. 6e,d** and **Supplementary Fig. 14d**). Importantly, similar experiments in U2OS cells yielded analogous results (**Supplementary Fig. 15**), demonstrating that our findings are not limited to a single cell line.

**Figure 6.**
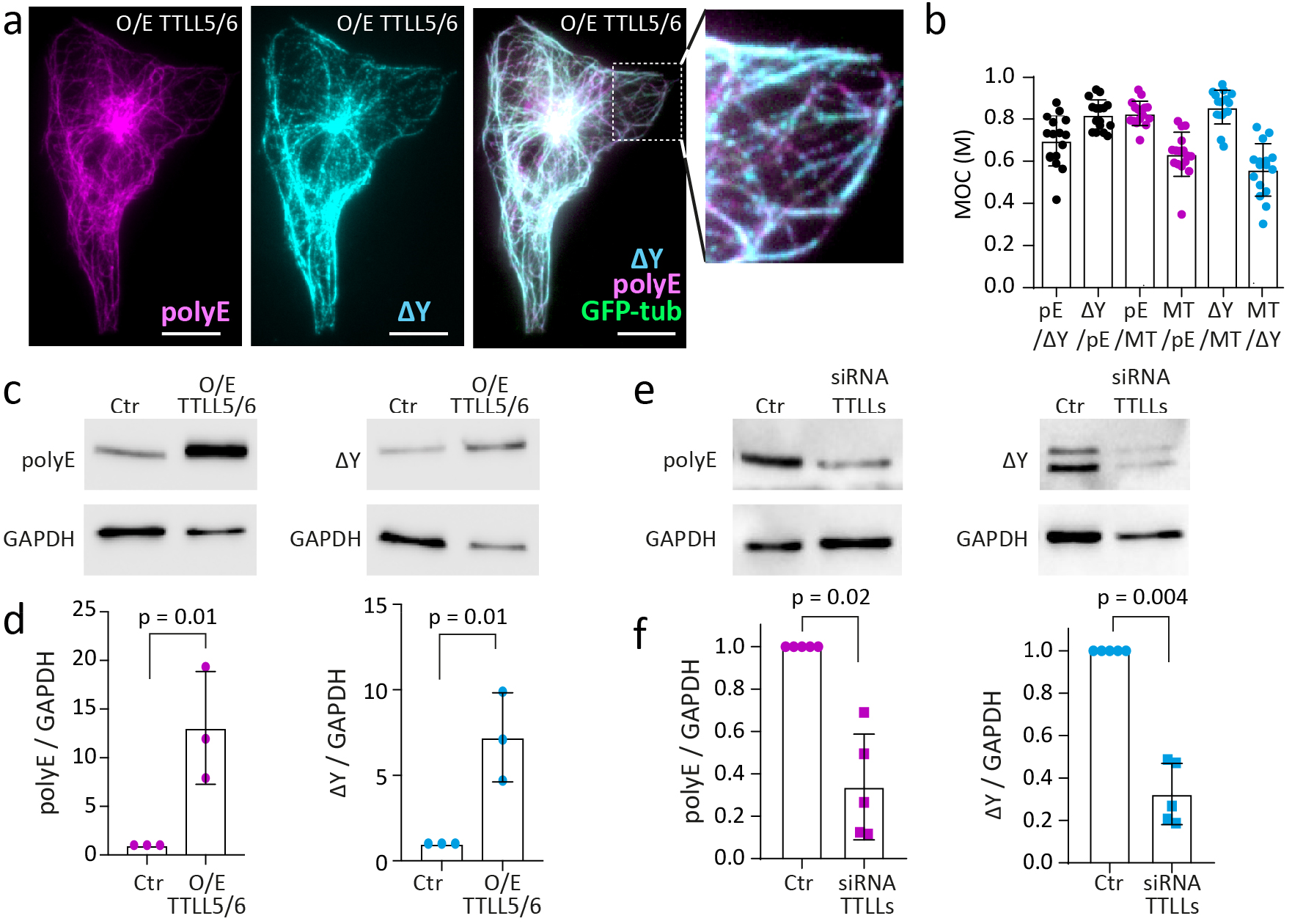
Detyrosination is determined by polyglutamylation in RPE-1 cells. **a**) Immunostaining of RPE-1 cells overexpressing TTLL5/6 (GFP: α-tubulin, magenta: polyE, cyan: ΔY). Boxed region is shown magnified on the right. **b**) Quantification of colocalization of polyE and ΔY via Mander’s overlap coefficient (MOC), measured for RPE1 cells overexpressing TTLL5/6. Mean with SD for 3 individual experiments of a total of N = 30 cells. **c**) Representative Western blot with **d)** quantification of tubulin PTM levels of the indicated control RPE1 cells (Ctrl) and RPE1 cells overexpressing TTLL5/6 (O/E TTLL5/6) using indicated antibodies. GAPDH antibody was used as loading control. Mean with SD of 3 independent experiments. Statistics: ANOVA, two-factor without replication. **e**) Representative Western blot with **f)** quantification of tubulin PTM levels in control siRNA (Ctrl) RPE1 cells and after knockdown of TTLLs (1, 5, 9, 11, 13) (siRNA TTLLs) in RPE1 cells using indicated antibodies. GAPDH antibody was used as loading control. Mean with SD of 5 independent experiments. Statistics: ANOVA, two-factor without replication. For all repeats, see **Supplementary Fig. 14**.

Together, our combined *in vitro* and in cell results support a model where polyglutamylation on microtubules controls local detyrosination by directing vasohibin/SVBP activity. This reveals a direct crosstalk between these two important tubulin PTMs, demonstrating that polyglutamylation primes microtubules for detyrosination.

## Discussion

In organisms, tubulin CTTs form a critical regulatory hub of microtubule function, with perturbations of PTMs leading to neurodegeneration^47^, cilia/flagella dysfunction and subfertility^48^ or loss of centriole integrity^49^. The development of recombinant tubulin expression protocols^22^ allowed to study *in vitro* how tubulin isotypes control motor activity^21^, or change microtubule lattice organization, protofilament numbers as well as microtubule stability^16,50^. Recombinant methods also enabled to introduce detyrosination and other proteolytic modifications of the tubulin tail, revealing changes in interactions with microtubule-associated proteins^51^. However, complete chemical control over the PTM patterns of tubulin CTTs was not available.

Here, we report the development of a modular, efficient and robust semisynthetic procedure to generate tubulins containing defined PTMs within their CTTs, using a split-intein based tandem transamidation approach^24^. Our method now allows to introduce any combination of PTMs via peptide synthesis, with full control over their type, chemical linkage, or position. Moreover, our approach is modular, as recombinant tubulins carrying a TTP-tag can be readily reacted with libraries of differently modified peptides. Semisynthetic designer tubulins thus provide a novel avenue to dissect the functional interplay between modified microtubules, MAPs and tubulin modifying enzymes in different systems.

Using our semisynthetic tubulin, we dissected PTM crosstalk between polyglutamylation and detyrosination on α-tubulin. Both PTMs co-localize on long-lived microtubules in neurons and centrioles, as well as on dynamic structures such as mitotic spindles^52^. We thus hypothesized that a crosstalk mechanism exists between both polyglutamylation and detyrosination. Using semisynthetic microtubules, containing either no PTMs, or glutamate branch chains of five, or ten glutamates at E445, we demonstrated that the activity of the enzyme responsible for tubulin detyrosination, vasohibin/SVBP, occurs in a polyglutamate branch length-dependent manner. Polyglutamylation of increasing chain length might either strengthen vasohibin/SVBP tubulin tail-interactions, or increased multivalency and charge density on the microtubule surface may enhance vasohibin diffusion on the microtubule surface^53^. While further studies will be required to determine the underlying biophysical mechanisms, our studies reveal that vasohibin mediates a direct crosstalk between polyglutamylation and detyrosination. This crosstalk might also work in both directions, as recent studies indicated that the activity of TTLL6, which elongates polyglutamate chains, is increased on detyrosinated tubulin peptides and microtubules^54^. Understanding the biological relevance of the coupling between polyglutamylation and detyrosination will be an exciting avenue for future research.

Our methods to semi-synthetically produce modified tubulin proteins are suitable to uncover and dissect further PTM interactions via writer and eraser enzymes in the future and allow to characterize the effect of such PTMs on both intrinsic microtubule properties and on reader proteins to gain a detailed understanding of the tubulin code.

## Acknowledgements

We thank Joel Schaer, Karsten Kruse, Emmanuel Derivery and Gina Grammbitter for discussions and comments on the manuscript. We thank Bertrand Beckert and Sergey Nazarov of Dubochet Center for Imaging for help with Cryo-EM data acquisition. We would like to thank Jijumon A.S. and Satish Bodakuntla (Institut Curie) for technical support. SF, MCV and CA have been supported by the DIP of the Canton of Geneva, SNSF (31003A_182473), and the NCCR Chemical Biology. BF and PG thanks the NCCR Chemical Biology and EPFL for support. CJ is supported by the French National Research Agency (ANR) award ANR-17-CE13-0021 and the Fondation pour la Recherche Medicale (FRM) grant DEQ20170336756.

## Author contributions

BF conceived the project. BF, PG and CA designed the experiments. EE and BF developed the tubulin semi-synthesis strategy with input from PG, GH and CA. EE performed peptide chemistry, tubulin expression and semi-synthesis, enzymatic assays and microtubule imaging, under supervision from BF. NA, NG, TR, LR and CJ contributed reagents for tubulin expression and semi-synthesis. SF and MCV performed cell experiments, microtubule imaging and analysis of *in vitro* tubulin dynamics, under supervision of CA. GH and FS performed cryoEM analysis, under supervision of PG. BF and CA initially wrote the manuscript and all authors contributed to data analysis, discussion of results and editing of the manuscript.

